# GEDI: an R package for integration of transcriptomic data from multiple high-throughput platforms

**DOI:** 10.1101/2021.11.11.468093

**Authors:** Mathias N. Stokholm, Maria B. Rabaglino, Haja N. Kadarmideen

## Abstract

Transcriptomic data is often expensive and difficult to generate in large cohorts in comparison to genomic data and therefore is often important to integrate multiple transcriptomic datasets from both microarray and next generation sequencing (NGS) based transcriptomic data across similar experiments or clinical trials to improve analytical power and discovery of novel transcripts and genes. However, transcriptomic data integration presents a few challenges including re-annotation and batch effect removal. We developed the Gene Expression Data Integration (*GEDI)* R package to enable transcriptomic data integration by combining already existing R packages. With just four functions, the *GEDI* R package makes constructing a transcriptomic data integration pipeline straightforward. Together, the functions overcome the complications in transcriptomic data integration by automatically re-annotating the data and removing the batch effect. The removal of the batch effect is verified with Principal Component Analysis and the data integration is verified using a logistic regression model with forward stepwise feature selection. To demonstrate the functionalities of the *GEDI* package, we integrated five bovine endometrial transcriptomic datasets from the NCBI Gene Expression Omnibus. The datasets included Affymetrix, Agilent and RNA-sequencing data. Furthermore, we compared the *GEDI* package to already existing tools and found that *GEDI* is the only tool that provides a full transcriptomic data integration pipeline including verification of both batch effect removal and data integration.

## Introduction

In gene expression data analysis, a higher number of samples leads to more statistically reliable results. Transcriptomic data integration is a useful computational tool to increase the sample size by combining multiple gene expression datasets. However, there are a few challenges when integrating transcriptomic data. These challenges include ID mapping, batch effect and how to merge microarray data with RNA-sequencing (RNAseq) data.

Microarray data are annotated with probe IDs that differ depending on the microarray platform, and RNAseq data are annotated with read IDs. Here we refer to both probe IDs and read IDs as reporter IDs. All the different types of reporter IDs must be mapped to the same ID type to integrate the datasets, which is possibly the biggest challenge in transcriptomic data integration.

The batch effect is the variance between datasets caused by different experimental conditions. This variance makes it impossible to draw any meaningful conclusions from the data. Therefore, the batch effect must be removed from the integrated dataset.

Even if the datasets have been re-annotated and a method to remove the batch effect is prepared, it is still not possible to integrate microarray data with raw RNAseq count data. Microarray data describes the relative expression profile compared to a reference sample whereas RNAseq data represents the expression profile with read counts. Therefore, before integration, RNAseq counts needs to be transformed to get the same type of numerical data as microarray data.

To the best of our knowledge, no current tools provide a transcriptomic data integration pipeline that addresses all the before-mentioned challenges. However, there are multiple tools that addresses some of the challenges. We will review some of them here.

COMMAND>_ (Moretto et al., 2019) is a web-based tool to search and download gene expression data directly from databases and map probes to genes using the probe sequences. One of its strengths is that no programming experience is required. Another strength is that COMMAND>_ is a multi-user tool which is ideal for teamwork. However, one of its weaknesses is that COMMAND>_ doesn’t remove the batch effect in the integrated data. And since COMMAND>_ isn’t accessible by R, it cannot be implemented in the transcriptomic data integration pipeline described here.

BioMart is an annotation database that can be used to map reporter IDs to gene IDs. The user does not have to know the probe sequences but can just use the probe IDs from the data table. The database can be used in a browser or with the *biomaRt* R package (Durinck et al., 2005, 2009).

The *sva* R package (Leek et al., 2020) is a tool that among other things can remove the batch effect from both microarray and RNAseq data using the functions ComBat and ComBat_seq respectively. Other recognized transcriptomic data analysis packages like *BingleSeq* (Dimitrov and Gu, 2020) and *tidybulk* (Mangiola et al., 2021) uses the *sva* package to remove batch effect. One downside to the ComBat functions is that they ignore genes with zero variance or zero counts within any batch. This can result in an unsuccessful batch correction if the variance between the batches is sufficiently large which can be the case when integrating data from different experiments and data types.

We have developed the Gene Expression Data Integration (*GEDI*) R package that solves all the above mentioned challenges by implementing already existing R packages to read, re-annotate and merge the transcriptomic datasets after which the batch effect is removed, and the integration is verified (Figure 1). The *GEDI* package addresses all the major challenges in transcriptomic data integration, unlike other solutions mentioned earlier. Therefore, we believe that the *GEDI* R package is a novel tool for transcriptomic data integration.

**Figure 1:**
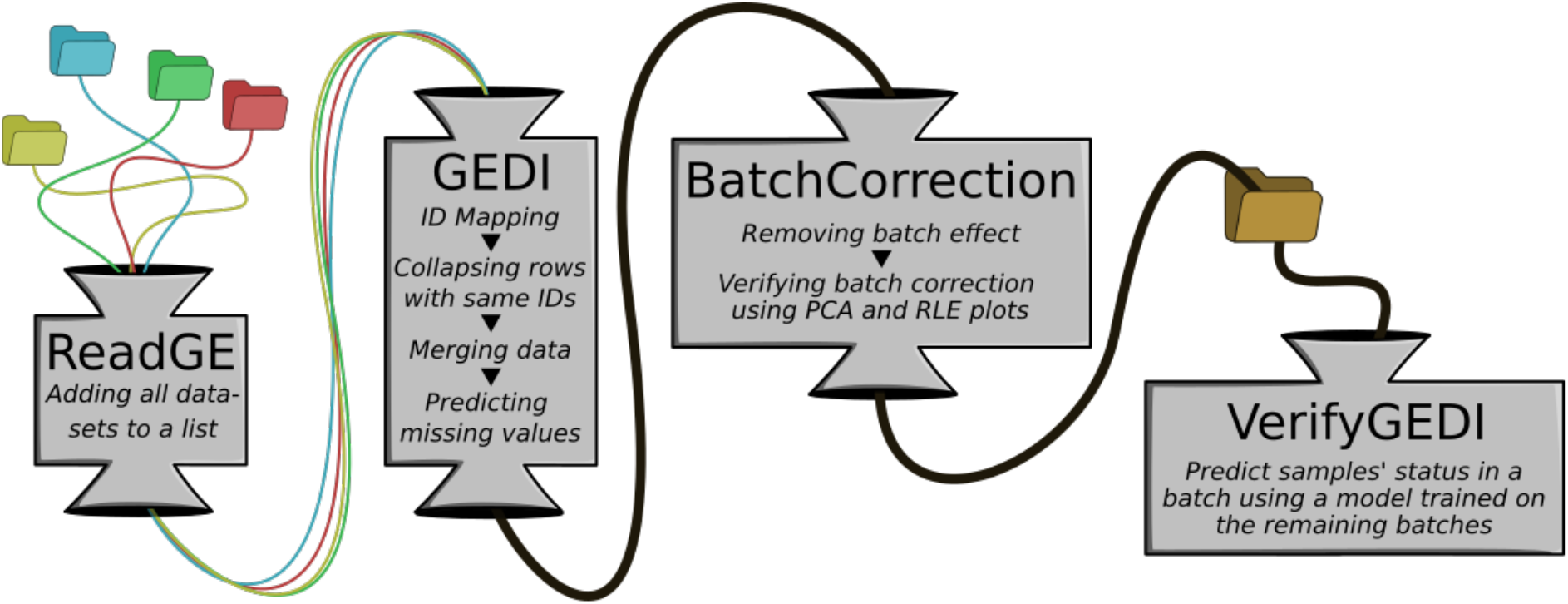
Flow diagram of the transcriptomic data integration pipeline provided by the GEDI package. First, the transcriptomic datasets, represented by colored folders, are read using the ReadGE function. The GEDI function integrates these datasets. This results in one transcriptomic dataset annotated with Ensembl or Entrez gene IDs. In the next step, the batch effect is removed by the BatchCorrection function, and it is verified with a Principal Component Analysis (PCA) plot and an RLE plot. Finally, the VerifyGEDI function verifies the data integration using a logistic regression model.

The entire *GEDI* workflow has been packaged into only four functions that allow the user to write a transcriptomic data integration pipeline in only 10-20 lines of code. The first function, ReadGE, reads the datasets and stores them in a list. It can read data from RNAseq and Affymetrix- and Agilent microarrays. The second function, GEDI, maps reporter IDs to Ensembl or Entrez gene IDs and merges the datasets into a single data table. The third function, BatchCorrection, removes the batch effect using modified functions from the *sva* R package (Leek et al., 2020). The batch correction is verified using Principal Component Analysis (PCA) and Relative Log Expression (RLE) plots. The fourth and final function, VerifyGEDI, verifies the transcriptomic data integration using a logistic regression model. These functions were demonstrated on a case inspired by a transcriptomic integration study by (Rabaglino and Kadarmideen, 2020).

## Method

### ReadGE Function

The ReadGE function reads multiple gene expression datasets and stores them in a list. The *GEDI* package can integrate three types of transcriptomic data: Affymetrix, Agilent and RNAseq. Affymetrix data is read using the *gcrma* R package (Wu and with contributions from James MacDonald Jeff Gentry, 2020), Agilent with the *limma* R package (Ritchie et al., 2015) and RNAseq with the *DESeq2* R package (Love et al., 2014).

One of the challenges mentioned in the Introduction was to merge microarray and RNAseq data. To address this challenge, the ReadGE function will transform RNAseq counts using variance-stabilizing transformation from the *DESeq2* package to yield the same type of numerical data as microarray data. Because ReadGE transforms the data, it is important that the RNAseq data consists of raw read counts. The counts are not transformed if only RNAseq data is integrated.

### GEDI Function

The GEDI function integrates transcriptomic datasets into one data table containing samples from all the datasets. This is done in three steps: ID mapping, collapse rows and merge datasets.

The ID mapping is done using the *biomaRt* R package (Durinck et al., 2005, 2009). Here the reporter IDs are mapped to either Ensembl or Entrez gene IDs. In case BioMart doesn’t contain annotation data from one of the platforms used, the user must provide an annotation table with reporter IDs in the first column and the corresponding Ensembl or Entrez gene IDs in the second column. See Table 2 for an example.

**Table 1:** Overview of transcriptomic datasets used for the case study. Each dataset is treated as a batch. There are three Affymetrix, one Agilent and one RNAseq dataset. The status is a variable describing if the endometrial sample is receptive (R) or non-receptive (nonR). The table shows how many R and nonR samples there are for each dataset.

| GEO Accession Number | Batch | Data Type | Status | Authors and year of publication |
| --- | --- | --- | --- | --- |
| GSE29853 | A | Affymetrix | R=6<br>nonR=6 | (Killeen et al., 2014) |
| GSE115756 | B | RNAseq | R=8<br>nonR=9 | (Rabaglino and Kadarmideen, 2020) |
| GSE107741 | C | Agilent | R=6<br>nonR=5 | (Matsuyama et al., 2018, *Article not published*) |
| GSE36080 | D | Affymetrix | R=3<br>nonR=3 | (Ponsuksili et al., 2012) |
| GSE20974 | E | Affymetrix | R=3<br>nonR=3 | (Salilew-Wondim et al., 2010) |

**Table 2:** First 4 rows of the annotation table for GSE107741. Can be used to re-annotate data if BioMart doesn’t contain the reporter IDs. First column is the reporter IDs of the dataset and the second column is the corresponding Ensembl or Entrez gene IDs.

| Reporter ID | Ensembl gene ID |
| --- | --- |
| A_73_112733 | ENSBTAG00000000005 |
| A_73_109892 | ENSBTAG00000000008 |
| A_73_P066806 | ENSBTAG00000000009 |
| A_73_113020 | ENSBTAG00000000010 |
| ... | ... |

Multiple reporter IDs may map to the same gene ID. Therefore, all rows with identical gene IDs are collapsed to go from probe-level to gene-level data. This is done using the *WGCNA* R package (Miller et al., 2011). The default method to collapse rows is to select the row with the highest mean and remove the rest.

The last step is to simply merge the datasets. Optionally, the GEDI function can predict missing expression values for genes where the percentage of known expression values exceeds a certain threshold defined by the user. The expression values are predicted using a linear regression model with one feature: the gene most correlated with the gene with missing values. The gene used as the feature for the model cannot have any missing values. Only one feature is chosen to get a quick estimate.

### BatchCorrection

The BatchCorrection function removes the batch effect from the integrated dataset. This function is inspired by the ComBat and ComBat_seq functions from the *sva* R package (Leek et al., 2020). These functions do not remove batch effect in genes with zero variance or zero counts within any batch. If these genes are ignored, there will still be variance between batches and the batch effect will remain. Therefore, the BatchCorrection function is modified to remove the batch effect in all genes.

The BatchCorrection function verifies itself visually with a before/after comparison of Principal Component Analysis plots and RLE plots. If the user chooses, boxplots can be used instead of RLE plots. To support this, a numeric output of the means and standard deviations of the gene expression values for each batch before and after the batch correction.

In a dataset with batch effect, the samples will aggregate in batches in the PCA plot, and the distribution is uneven in the RLE plot. The mean expression values will also differ in the different batches. If the batch effect is successfully removed, the samples will no longer aggregate in batches and the all the samples will have a similar distribution. The mean expression values should also be similar. An example of the visual and numeric output can be seen in Figure 2 and Table 3 respectively.

**Table 3:** Numeric output from the BatchCorrection function. Mean (M) and standard deviation (SD) of the gene expression values are calculated for each batch/dataset before and after the batch correction. After a successful batch correction the means for all batches should be similar.

| Batch | Before Batch Correction | After Batch Correction |
| --- | --- | --- |
| A | M (SD) = 7.44 (0.03) | M (SD) = 8.19 (0.03) |
| B | M (SD) = 8.35 (0.02) | M (SD) = 8.19 (0.02) |
| C | M (SD) = 9.97 (0.02) | M (SD) = 8.19 (0.02) |
| D | M (SD) = 6.92 (0.05) | M (SD) = 8.19 (0.04) |
| E | M (SD) = 7.23 (0.02) | M (SD) = 8.19 (0.01) |

**Figure 2:**
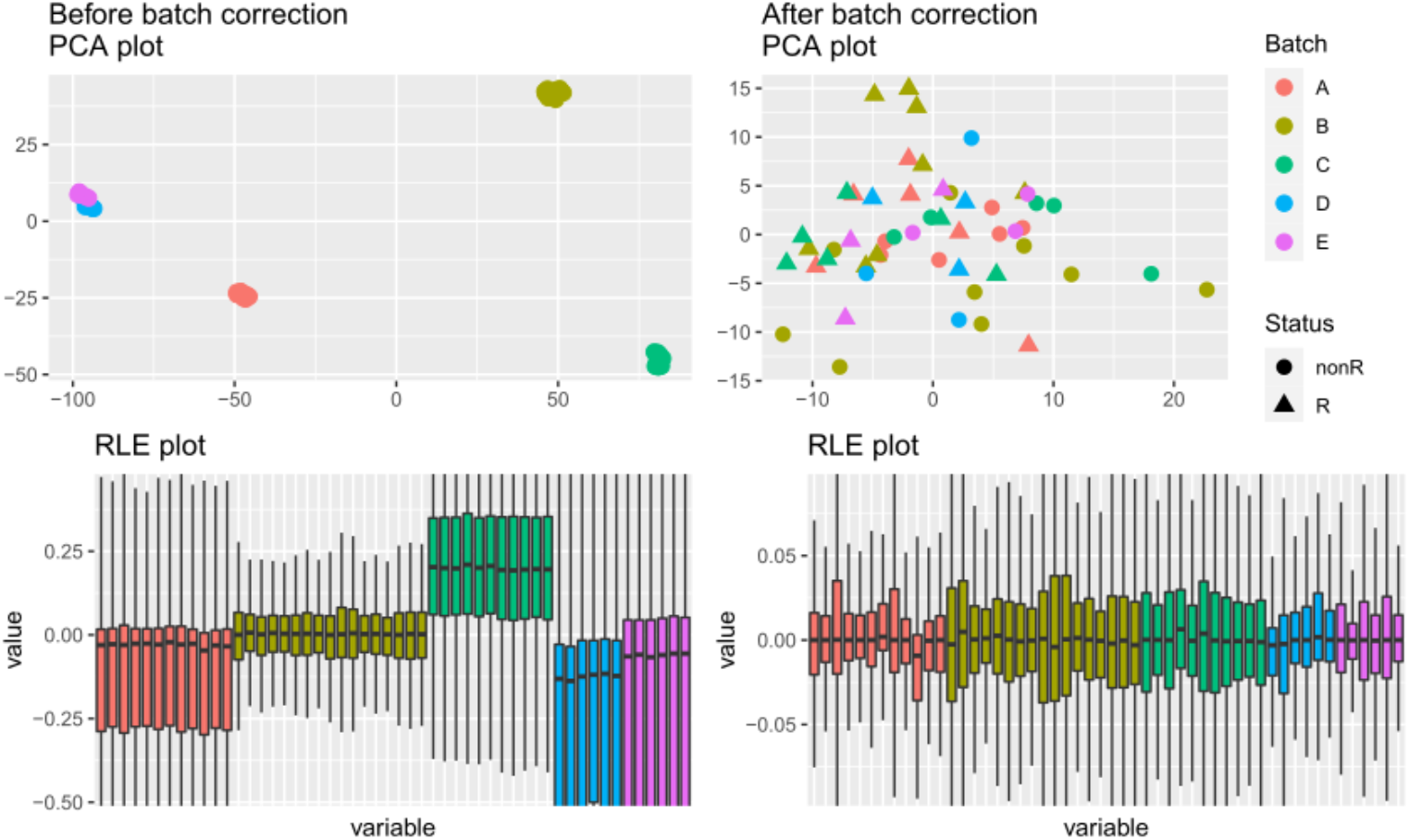
The PCA and RLE plots are created by the BatchCorrection function to verify batch effect removal. Each color represents a batch/dataset and the shapes in PCA plot corresponds to the status of the samples. Before the batch correction, the samples aggregate in batches in the PCA plot, and the distribution is uneven in the RLE plot. After a successful batch correction, the samples no longer aggregate in batches and all samples have as similar distribution.

### VerifyGEDI

The VerifyGEDI function verifies the transcriptomic data integration using a logistic regression model to predict the status of the samples in one batch based on the remaining batches’ samples. Here the status of a sample refers to some parameter like sick/healthy or treated/control. If the accuracy of this model is high, then it must mean that the batches have similar data and biological background. Therefore, the data integration can be verified using this method.

The default model is a logistic regression model and the forward stepwise selection (FSS) algorithm picks which genes to use as features. The accuracy of the model is measured using cross-validation so each of the datasets are used as test-data exactly once. The function output shows the features in the final model as well as its accuracy. If the accuracy is significantly high, the data integration was successful.

### Case study

The functionalities of the *GEDI* package were tested using data from a study by (Rabaglino and Kadarmideen, 2020) where five endometrial transcriptomic datasets from cattle were integrated. The datasets were downloaded from the NCBI Gene Expression Omnibus (GEO) with the following accession numbers GSE29853, GSE115756, GSE107741, GSE36080 and GSE20974. An overview of the five datasets used can be seen in Table 1. This case was ideal to demonstrate the *GEDI* package since Agilent, Affymetrix and RNAseq datasets were integrated.

## Results

### Case study

Using the ReadGE function, the datasets were read and stored in a list without any issues. The second step in the pipeline is to use the GEDI function. The Agilent dataset (GSE107741) is annotated with probe IDs that does not exist in BioMart. Therefore, an annotation table is created using the platform information found in GEO. Table 2 shows the first 4 rows of the annotation table for GSE107741. We chose to predict missing expression values with a threshold of 75%. This resulted in expression values being estimated for 1877 genes. This means that the dataset kept 1877 genes that would otherwise have been removed. However, up to 25% of the expression values in those genes are estimated by a linear regression model.

Next, the batch effect is removed using BatchCorrection. The visual output of the function can be seen in Figure 2. Here, the data is aggregated in batches in the PCA plot before the batch correction, and the data distribution is uneven in the RLE plot. After the batch correction, the data no longer aggregate in clusters, and all the samples have a similar distribution. This indicates that the batch correction was successful. The numeric output supports this conclusion since the mean expression values are equal in all batches after the batch correction (Table 3).

As the final step in the pipeline, VerifyGEDI is used to verify the data integration with a logistic regression model. The forward stepwise selection (FSS) algorithm chose 4 genes as features for the model: ENSBTAG00000013072, ENSBTAG00000000476, ENSBTAG00000003826 and ENSBTAG00000000188. The cross-validation showed an accuracy of 91.7%. This high accuracy leads to conclude that the transcriptomic data integration was successful. Furthermore, the four selected genes are potential biomarkers for endometrial receptivity in cattle.

## Discussion

In studies involving feature or target discovery for biotechnological or biomedical fields, it is often necessary to integrate transcriptomic datasets from both spatial and temporal environments. Here, *GEDI*, an R package, was developed to allow users to quickly write a transcriptomic data integration pipeline that besides integrating the datasets, also verifies that the integration was successful and that the batch effect has been removed.

The current tools we have reviewed here, provide only part of the transcriptomic data integration pipeline. Some tools like COMMAND>_ (Moretto et al., 2019) re-annotate and integrates the datasets, but the batch effect is not removed. The *sva* R package (Leek et al., 2020) removes the batch effect, but genes with zero variance or zero counts within any batch are not corrected. Because of this, it is not always sufficient to simply use the *sva* R package after integrating data in COMMAND>_. And manually combining multiple tools to integrate data is not convenient for the user.

Besides all its strengths, *GEDI* has some limitations. *GEDI* can only integrate three types of datasets: Affymetrix, Agilent and RNAseq. If the user wants to use other types of gene expression datasets, then the datasets must be read and stored in a list without using the ReadGE function. Another weakness is the dependency on the *biomaRt* R package (Durinck et al., 2005, 2009). When *GEDI* maps reporter IDs to Ensembl or Entrez gene IDs, the Ensembl site used by the *biomaRt* package can be unresponsive. The only solution to this is try again later or use annotation tables as shown in Table 2. At the time of publication, the *GEDI* package can only map to either Ensembl gene IDs or Entrez gene IDs. Other than limitations for integration, verification of integration is limited to data where a status variable is known for all datasets. Meaning the datasets must have a similar biological background. This limitation is intentional since the *GEDI* package is not meant to be used for unsupervised analysis except for when verifying the batch correction.

The case study demonstrated the functionalities of the *GEDI* package and showed that it successfully addressed all the challenges mentioned in the Introduction. RNAseq data was integrated with microarray data, the datasets were re-annotated using *biomaRt* and the batch effect was removed using modified ComBat and ComBat_seq functions from the *sva* package.

*GEDI* facilitates a straightforward integration of any number of transcriptomic datasets, and it can be applied in studies working with multi-transcriptomic data from any species. The package provides a full transcriptomic data integration pipeline including verification of both batch correction and data integration. Future development of *GEDI* may include allowing the ReadGE function to automatically download transcriptomic datasets from databases like GEO.

## Data Availability Statement

The datasets used in the Results section were downloaded from the open-source database GEO. The *GEDI* package (licence: CC BY-NC-SA 4.0) can be installed from https://github.com/MNStokholm/GEDI. Package updates after publication will also be uploaded to GitHub. An example pipeline and data can be found in GEDI/man/Examples in the GitHub repository. Further inquiries can be directed to the corresponding author.

## Authors’ contribution

MNS developed the computational architecture and scripts used to develop this GEDI pipeline under the guidance of MBR and HNK. MNS wrote the first draft. HNK and MBR conceived the project and improved various drafts of this manuscript. All authors read and approved the final version.

## Ethical Statement

Not Applicable

